# Identifying human-specific transcription factor binding site gains and losses that contribute to human-specific phenotypes

**DOI:** 10.64898/2026.09.06.747930

**Authors:** Manuel Ferrando-Bernal, John A. Capra

## Abstract

Identifying the genetic changes responsible for human-specific traits remains a central challenge in evolutionary genomics. Although thousands of modern human-specific variants have been identified through comparisons with Neanderthal, Denisovan, and great ape genomes, the phenotypic consequences of most of these changes remain unknown, particularly for non-coding variants. Here, we predicted transcription factor binding site (TFBS) gains and losses for 16,883 human-specific fixed and high frequency variants. We identified 3,357 human-specific variants within cis-regulatory elements (cCREs) that modify TFBS motifs and prioritized variants for which both the target gene and the affected transcription factor are associated with the same phenotype. This identified 404 candidate human-specific regulatory variants involving 387 genes. These genes are enriched for skeletal traits known to differ between modern humans and archaic hominins. Beyond skeletal morphology, enrichment analyses suggested that human-specific regulatory changes also affect soft tissues that are not represented in the fossil record, including behavioral, reproductive, and facial traits. Independent support for the functional relevance of these candidates comes from their significant enrichment in modern human-derived differentially methylated regions and among variants experimentally shown to alter gene regulatory activity in massively parallel reporter assays. Furthermore, candidate variants occur within unusually large regions depleted of archaic introgression, consistent with the hypothesis that some regulatory changes contributed to reduced fitness following admixture with Neanderthals and Denisovans. Together, our results provide a prioritized catalogue of human-specific regulatory variants and reveal candidate molecular mechanisms underlying traits that distinguish modern humans from archaic hominins.

## Introduction

Understanding the genomic changes behind human-specific traits is a main goal of the study of human evolution, but the genetic basis for humanness remains largely unknown. The availability of high-coverage archaic genomes from Neanderthals and Denisovans has enabled the characterization of thousands of alleles that are fixed or nearly fixed in modern humans but absent in the other hominids (1-3). While some of these changes affect protein-coding sequences (4-8), the vast majority are in non-coding regions, suggesting that the changes in gene regulation play a major role in the evolution of human-specific traits (3).

Modification of transcription factor binding sites (TFBSs) is a driver of gene regulatory evolution. One of the first functional studies of human-specific non-coding variants examined 25 TFBSs with human-specific alleles (9). Using reporter assays in neuronal cell lines, they found that half of the human variants altered gene expression compared to the ancestral allele. Since this analysis, several additional high archaic genomes have been sequenced (10-13), as well as several additional species of non-human primates. Our knowledge of cis-regulatory regions (cCREs) and TFBS has also expanded substantially (14, 15). This facilitates the identification of alleles that are specific to modern humans (1, 3) and opens an opportunity to explore how human-specific genomic changes affect cis-regulatory activity to influence human-specific traits. More recently, genome-wide analysis using massively parallel reporter assays (MPRA) demonstrated that hundreds of human-specific alleles have cis-regulatory activity in neurons, osteoblasts, and embryonic stem cells, and that these alleles are enriched for divergent TFBS motifs (3). However, the contexts in which human-specific variants have gene regulatory functions and whether these changes contribute to specific phenotypes that distinguish modern humans from archaic hominins remains poorly understood.

Here, we combined two recent studies that identified variants derived in the modern human lineage after comparing the genomes of thousands of modern humans and several high-coverage Neanderthals, Denisovans, and four non-human primate species (1, 3). Together, these datasets yielded approximately 17,000 candidate modern human-specific high frequency variants. Based on the expectation that non-coding changes mediating TF binding account for a substantial fraction of phenotypic differences between closely related species, we focused on variants capable of creating or disrupting novel TFBS in known cCREs. We then evaluated the contribution of these alleles to the evolution of modern traits since the divergence with the ancestors of Neanderthals and Denisovans. Together, our results identify TFBS motif changes in cCREs that likely contributed to the emergence of many human-specific traits, giving new evidence of how regulatory variants play an important role in speciation.

## Results

### Identification of human-specific gains and losses of transcription factor binding motifs

In order to identify human-specific variants, we integrated results from two studies that compared present-day human genomes from the 1000 Genomes Project with high-coverage Neanderthal and Denisovan genomes and four great ape species (1, 3). These studies identified variants that are derived and fixed or nearly fixed in present-day humans while the ancestral allele is present in archaic hominins and other great apes. After merging both datasets, removing redundancies, and removing allelic mismatches, we obtained a set of 16,883 fixed or nearly fixed variants that we refer to as “human-specific” (Figure 1A).

**Figure 1.**
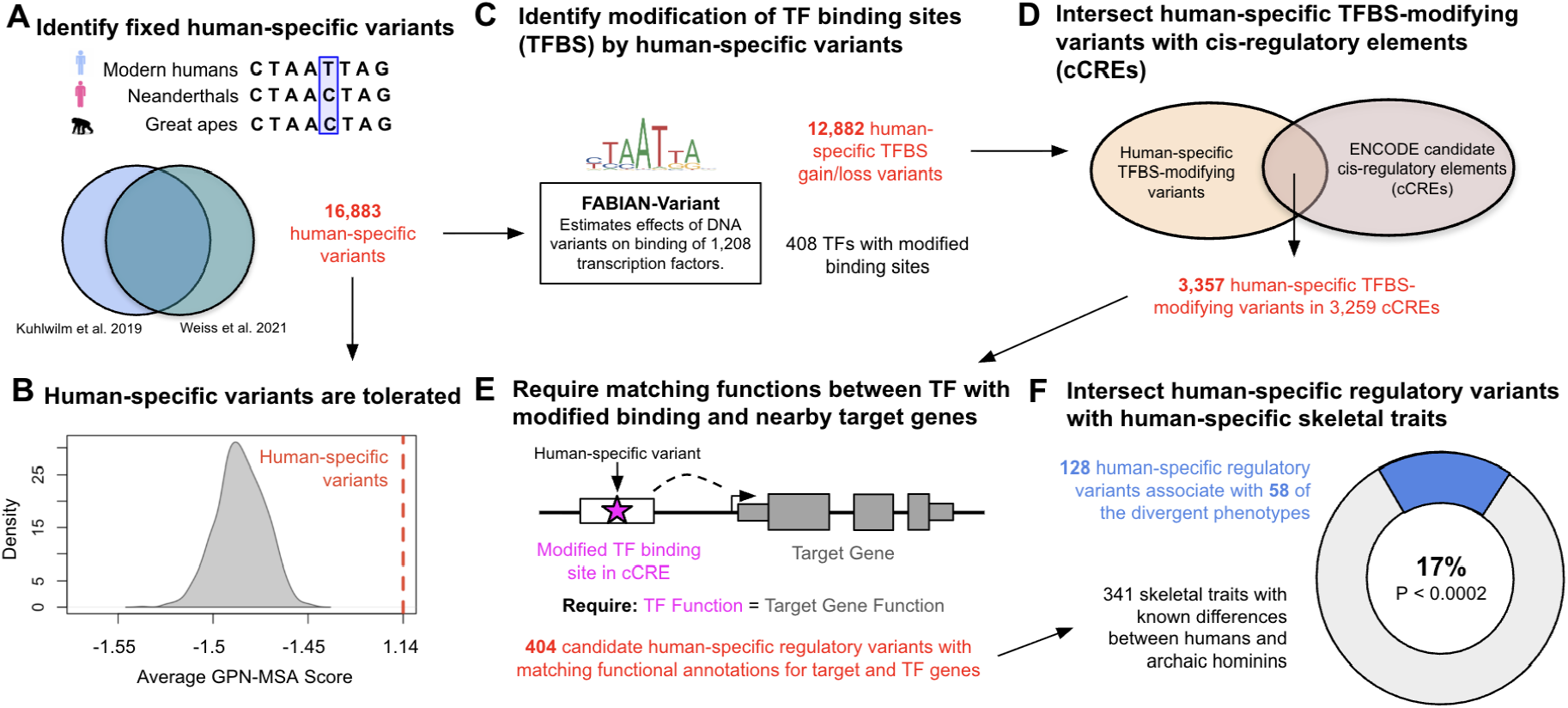
Framework for detecting candidate human-specific gene regulatory variants underlying phenotypic differences between humans and archaic hominins. **(A)** Human-specific variants from Kuhlwilm et al. (2019) and Weiss et al. (2021) were combined and mismatching alleles removed, giving a final set of 16,883 derived variants fixed or nearly fixed in modern humans compared to archaic hominins and great apes. **(B)** These human-specific alleles were scored using the GPN-MSA language model (Supplementary Table 1). The observed score (red dashed line, 1.14) is significantly higher than expected from the null distribution (gray), consistent with the human-specific allele being functionally tolerated. **(C)** Human-specific variants overlapping transcription factor binding sites were identified and tested for binding site creation or disruption using FABIAN-Variant. This yielded 12,882 human-specific variants predicted to modify TFBS motifs. **(D)** Intersecting the variants with known cis-regulatory elements (promoters, enhancers, and other cCREs) from ENCODE narrowed this down to 3,357 variants located within 3,259 cCREs. **(E)** Candidate regulatory variants were prioritized when both a nearby target gene and the transcription factor whose binding site was modified were linked to the same HPO phenotype. This yielded 404 candidate human-specific regulatory variants linked to 387 genes (Supplementary Table 2). **(F)** Focusing on 341 skeletal phenotypes known to be different among humans and archaics yielded 128 human-specific variants linked to 155 genes associated with 58 human-specific skeletal traits (Supplementary Table 4). This represents significant enrichment for these traits (P < 0.0002). These variants highlight plausible regulatory mechanisms behind the trait difference.

We evaluated the likelihood of the human-specific variants being tolerated using the GPN-MSA DNA language model (16). GPN-MSA is trained on genomic multiple sequence alignments of diverse vertebrate species to estimate the probability of each possible base at each position in the human genome and compare it to the human reference allele. GPN-MSA scores were available for 16,369 of the variants (Supplementary Table 1). The human-specific alleles have significantly higher GPN-MSA scores than expected (Figure 1B; average 1.14, P < 0.0001), suggesting that they are tolerated in the human lineage. We note for these variants the derived allele is the human reference, so the positive score indicates that the archaic allele has higher probability according to the model.

We tested if human-specific variants created or disrupted a motif for 1,208 human transcription factors using FABIAN-variant (17). Considering a strict threshold, 12,882 human-specific variants create or disrupt at least one TFBS motif, corresponding to <0.05% of all tested variant-model pairs (Figure 1C). The number of TFs with at least one binding site created or disrupted is 408 (400 created and 403 disrupted). The top five transcription factors with the most common modified binding sites are GATA2 (473 times disrupted, 378 created), FOXG1 (272 disrupted, 252 created), POU2F1 (369 disrupted, 251 created), ZBTB7A (247 disrupted, 247 created), and POU2F2 (296 created, 217 disrupted).

Since matches to binding motifs may not be functional, we prioritized motif changes with functional potential by overlapping human-specific variants modifying TFBS with cCREs annotated by ENCODE across the human genome (18). This dataset catalogs genomic regions with functional genomic signatures suggesting regulatory functions, including enhancers, promoters, and other regulatory elements, across hundreds of cell types. Of the 12,882 human-specific variants modifying TFBS, 3,357 overlapped 3,259 known regulatory regions (Figure 1D).

Finally, we required that both the TF that binds the motif and the target gene should have overlapping functional annotations (Figure 1E). Applying this strict criterion to each of the 3,357 TFBS variants yielded 404 human-specific variants linked to 261 target genes and 148 TFs (387 total; Supplementary Table 2). These variants are associated via these genes with 586 HPO traits. While this filter is likely conservative, we wanted to enrich our candidate set for changes with the strongest functional support.

### Human-specific candidate regulatory variants are significantly enriched for associations with known human-specific skeletal phenotypes

Next, we sought to evaluate if these human-specific TFBS changes in cCREs contribute to phenotypic traits that distinguish modern humans from archaic hominins. First, we obtained a curated set of 341 skeletal traits from the Human Phenotype Ontology (HPO) (19) known to be divergent based on comparisons with modern humans and Neanderthal fossil remains (Supplementary Table 3) (20). Given that soft tissue does not fossilize, this set of skeletal traits represent the best sources of direct phenotypic comparisons between modern and archaics.

The human-specific variants we prioritized associate with 58 of the known divergent skeletal phenotypes, significantly more than expected (Figure 1F, P < 0.0002). Specifically, we identified 128 human-specific TFBS altering variants in which both genes are annotated with at least one skeletal trait that differs between humans and archaics. These variants are associated with a total of 155 genes, including 66 with TFBS disruptions and 96 target genes (Supplementary Table 4). If we relax the threshold and require only one of the target genes or TF to be related for a divergent skeletal phenotype, we recover 203 of the 341 divergent phenotypes (Supplementary Table 5). These analyses identify human-specific regulatory candidates for the development of skeletal traits that make us unique.

Since it is possible that the human-specific variants disrupted a TFBS leading to a loss of an archaic cCRE, we repeated the analysis filtering by known cCREs only when the human-specific variant creates a new TFBS, but not when it disrupts at TFBS. This identifies regulatory candidate genes (including target genes and TFs) linked to 72 of the known differential skeletal traits (Supplementary Table 6). Relaxing the requirement that the target gene and the TF both associate with the same phenotype, the candidate variants are associated with 215 out of the 341 skeletal phenotypes (Supplementary Table 7).

### Human-specific regulatory variants identify novel candidate human-specific phenotypes

Given the strong enrichment for known divergent skeletal traits among human-specific TFBS changes, we expanded the analysis to other phenotypes not conserved in the fossil record. Considering all 387 genes linked to the 404 candidate human-specific TFBS changes in cCREs for which a target gene and its associated TF coincided in their functional annotation yielded 586 HPO traits (Supplementary Table 8), including the 58 known divergent skeletal traits.

These genes were significantly enriched for many HPO traits beyond the skeletal system (21) (FDR < 0.05, Table 1; Supplementary Table 9). For example, behavioral traits (autism, odds ratio [OR]=3.53), hair phenotypes (hypopigmentation, OR=15.3), reproductive traits (penis size OR=2.89 and cryptorchidism OR=1.85), and heart defects (hypoplastic heart OR=8.49) were all enriched over the HPO background. As expected given the enrichment for known skeletal differences between humans and archaic hominins, we also identified enrichment for many skeletal traits individually, some of which are known to differ, such as broad forehead and abnormality of the chin, but also in facial traits that do not fossilize like the lips. Genes linked to conditions with autosomal dominant inheritance patterns were also significantly enriched (OR=2.28), suggesting that for many of the variants a single allele was sufficient for substantial functional effects.

**Table 1.** Phenotypes significantly enriched among the genes linked to candidate human-specific regulatory variants. See Supplementary Table 9 for full enrichment test results.

| Term | Odds Ratio | Gene Overlap | P-value | Adjusted P-value |
| --- | --- | --- | --- | --- |
| Autosomal dominant inheritance (HP:0000006) | 2.28 | 135/1059 | 1.47E-12 | 1.99E-09 |
| Autism (HP:0000717) | 3.53 | 20/94 | 1.14E-05 | 0.01 |
| Brachydactyly syndrome (HP:0001156) | 2.57 | 30/184 | 2.42E-05 | 0.01 |
| White forelock (HP:0002211) | 19.13 | 6/10 | 2.51E-05 | 0.01 |
| Patchy hypopigmentation of hair (HP:0011365) | 15.30 | 6/11 | 5.19E-05 | 0.01 |
| Vesicoureteral reflux (HP:0000076) | 3.80 | 15/66 | 6.46E-05 | 0.01 |
| Patent ductus arteriosus (HP:0001643) | 2.84 | 22/123 | 7.36E-05 | 0.01 |
| Hypoplastic left heart (HP:0004383) | 12.75 | 6/12 | 9.73E-05 | 0.02 |
| Cryptorchidism (HP:0000028) | 1.95 | 46/360 | 1.19E-04 | 0.02 |
| Everted lower lip vermillion (HP:0000232) | 3.52 | 15/70 | 1.32E-04 | 0.02 |
| Sprengel anomaly (HP:0000912) | 5.13 | 10/35 | 1.45E-04 | 0.02 |
| Broad forehead (HP:0000337) | 3.17 | 15/76 | 3.45E-04 | 0.04 |
| Micropenis (HP:0000054) | 2.89 | 17/93 | 3.70E-04 | 0.04 |
| Hypoplastic heart (HP:0001961) | 8.49 | 6/15 | 4.36E-04 | 0.04 |
| Heterochromia iridis (HP:0001100) | 8.49 | 6/15 | 4.36E-04 | 0.04 |
| Deeply set eye (HP:0000490) | 3.07 | 15/78 | 4.64E-04 | 0.04 |
| Synophrys (HP:0000664) | 3.92 | 11/47 | 4.67E-04 | 0.04 |
| Widely spaced teeth (HP:0000687) | 5.38 | 8/27 | 5.17E-04 | 0.04 |
| Proximal placement of thumb (HP:0009623) | 6.38 | 7/21 | 5.26E-04 | 0.04 |
| Deviation of the hallux (HP:0010051) | 4.61 | 9/34 | 5.84E-04 | 0.04 |
| Esophageal atresia (HP:0002032) | 10.60 | 5/11 | 6.75E-04 | 0.04 |
| Clinodactyly of the 5th finger (HP:0004209) | 2.16 | 26/183 | 8.03E-04 | 0.05 |

As a complementary analysis, we also tested these genes for enrichment in organs, anatomical systems, and specific body parts using Gene ORGANizer (22). Supporting the HPO results, we found significant enrichment (FDR < 0.05) for the lips (1.42x) and forehead (1.31x), and suggestive enrichment (FDR < 0.1) for the maxilla and jaws (1.2x, Figure 2A; Supplementary Table 10). The strongest significant enrichment was for the hypothalamus (1.82x), a deep brain structure that is challenging to study directly from fossil evidence.

**Figure 2.**
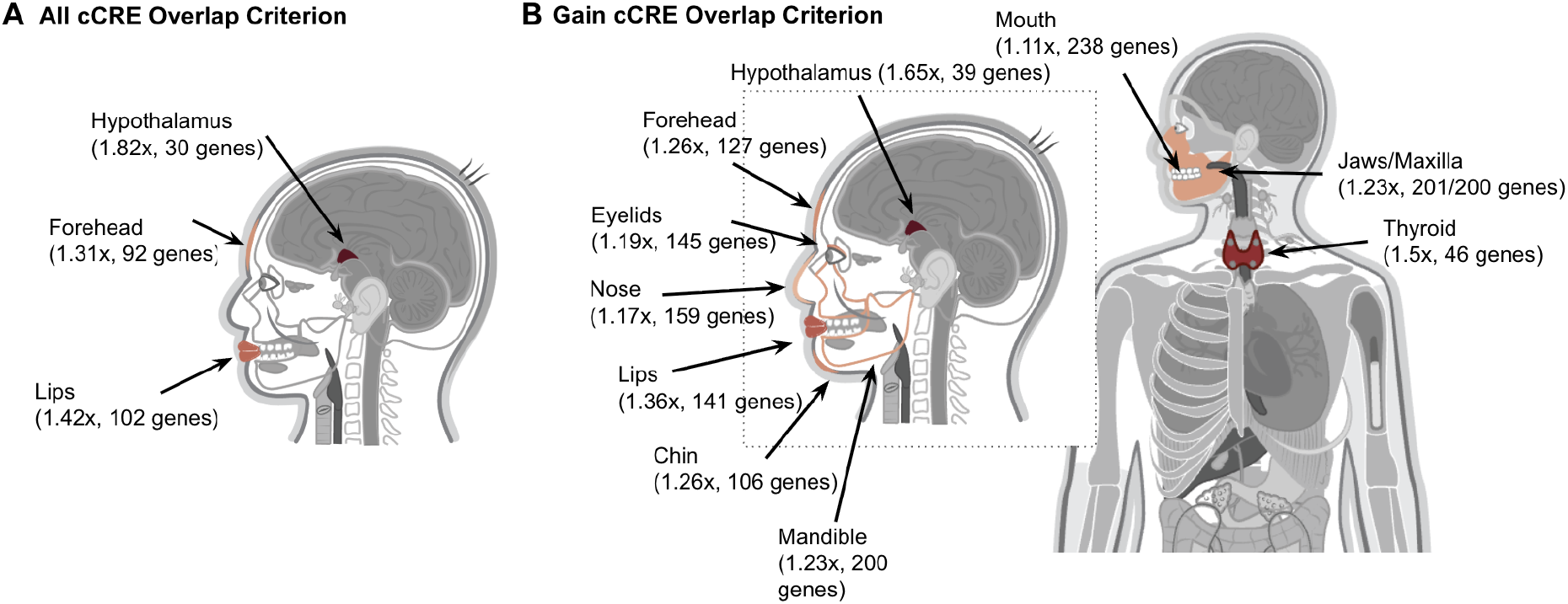
Organs with significant enrichment for candidate human-specific cis-regulatory variants. **(A)** The 387 candidate genes (261 target genes and 148 TFs) linked with human-specific regulatory variants are significantly enriched for functions in the lips, forehead, and hypothalamus compared to all genes in HPO (FDR < 0.05, geneORGANizer; Supplementary Table 10). Darker red indicates stronger enrichment. **(B)** Relaxing the requirement that TFBS losses overlap cCREs, we identified 579 candidate genes with significant enrichment for 11 anatomical structures, including the lips, jaw, maxilla, mandible, hypothalamus, forehead, thyroid, chin, eyelid, nose, and mouth (FDR < 0.05, geneORGANizer; Supplementary Table 11)).

Repeating this analysis without the requirement for TFBS losses to overlap cCREs identified 579 genes related to 893 traits in HPO where both the target gene and the TF share the same HPO trait and with enrichment for 11 organs in Gene ORGANizer, including the three highlighted above (hypothalamus, forehead, lips) and the jaw, maxilla, mandible, thyroid, chin, eyelid, nose, and mouth (Figure 2B; Supplementary Tables 11).

These results extend the enrichment observed for known skeletal differences to additional anatomical structures, including soft tissues that are not directly preserved in the fossil record, and suggest that human-specific regulatory changes may have contributed to species-specific differences in both skeletal and non-skeletal traits.

### Human-specific candidate regulatory variants are enriched for regions divergently methylated between humans and archaic hominins

To obtain further evidence for the functional relevance of candidate regulatory variants, we intersected them with previously published maps of divergent DNA methylation in bone cells between modern humans, archaic hominins and non-human primates (23). Of the 3,357 human-specific variants that altered TFBS in cCREs, 13 are within the 873 differentially methylated regions (DMRs) identified as derived in modern humans. While this is a small number due to the limited scope of the DMRs, it is significant enrichment relative to random expectation (P < 0.0022). Since the DMR boundaries are incomplete due to technical challenges of working with ancient DNA, we also considered 10,000 base pairs on either side. This identified 29 more DMRs (38 total, Supplementary Table 12) with 51 human-specific variants in cCREs modifying TFBS (P < 0.0001). This agrees with previous observations that human-specific variants tend to be closer to DMRs and may be responsible for some of them (23).

Two of the DMR variants were also among the 128 variants associated with human-specific skeletal phenotypes identified in our analyses, significantly more than expected by chance (P < 0.0217). Both are in regions that are hypermethylated in humans compared to the archaics and chimpanzees. One is located on chromosome 3 (chr3:71532882) in a DMR (chr3:71532285-71534235) in an intron of *FOXP1* (Figure 3). This position has a predicted change in a TFBS for *TFAP2B* modified by the human-specific derived allele. *TFAP2B* is involved in midfacial neural crest development (24), and *FOXP1* is known to be expressed during neurocranial development (25). These observations are consistent with both *FOXP1* and *TFAP2B* being related to malar flattening, a known differential trait among humans and archaics. This suggests that the human-specific genetic change led to differences in the regulation of *FOXP1* by TFAP2C ultimately contributing to this human-specific trait.

**Figure 3.**
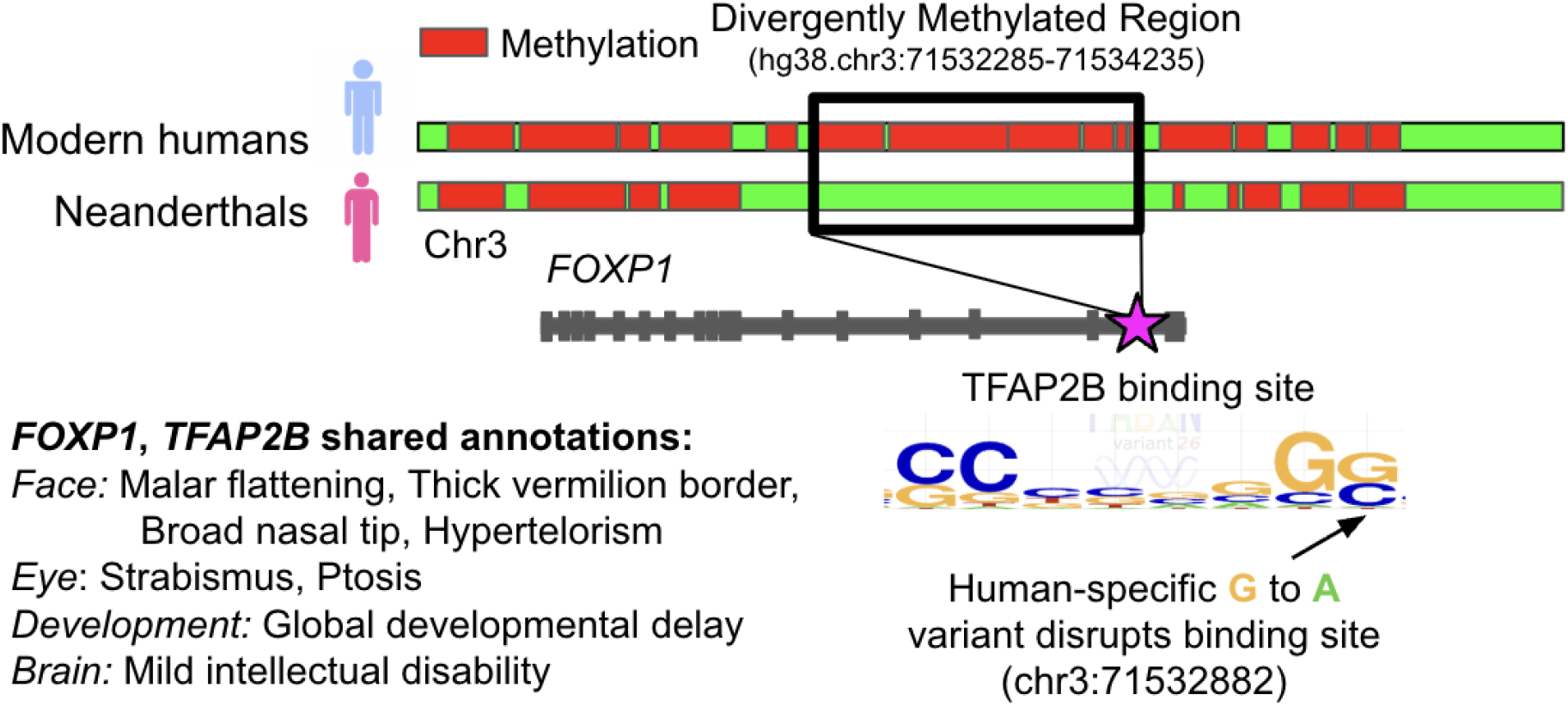
A human-specific variant disrupts a binding motif for TFAP2B in an enhancer in a hypermethylated region in an intron of FOXP1. Both genes are associated with several human-specific craniofacial phenotypes, as well as developmental delay and mild cognitive impairment.

The second variant (chr22:40327114) is located in a DMR (chr22:40326471–40331181) near *TNRC6B*. This variant creates a TFBS for *FLI1* in an annotated enhancer. Both *TNRC6B* (26) and *FLI1* (27) are highly expressed during embryonic development and both are linked to macrocephaly in HPO.

### Human-specific candidate regulatory variants are enriched for human-specific regulatory activity

We further evaluated the candidate human-specific TFBS changes using MPRA data measuring the regulatory consequences of human-specific variants. In a previous study, 407 human-specific variants were shown to produce significant expression differences between the modern human and archaic alleles based on an MPRA conducted on three different cell lines: neural progenitors, stem cell, and osteoblasts (3). These experimentally validated variants were significantly enriched among our candidates (21 variants out of our functional candidate set of 404 human-specific variants, P < 0.001, Supplementary Table 13), lending further support to their relevance to regulatory divergence. These validated human-specific regulatory variants affect TFs and target genes with clear connections to many divergent traits, including shallow orbits, mandibular prognathia, malar flattening, short nose, wide nasal bridge, and pectus excavatum. The 21 variants are related to 58 genes, and these genes are associated with 127 HPO phenotypes. These phenotypes are enriched for skeletal traits known to be different among humans and archaics (14 out the 127, P < 0.0001). For example, a human-specific allele within an intron of *FOXP1* (∼20 kb away from the DMR variant discussed above) alters a TFBS for *RUNX2*, an essential gene for craniofacial development (28). Both genes are associated with malar flattening and mandibular prognathia—skeletal phenotypes with morphological divergence between modern humans and archaic hominins.

Together, these independent lines of evidence support the functional relevance of the variants identified in our analysis. The enrichment of candidate variants within modern human-derived differentially methylated regions, combined with their overrepresentation among variants experimentally shown to alter gene expression, confirms that many human-specific mutations modifying TFBS in known regulatory elements have measurable regulatory consequences. These findings support the hypothesis that such variants contributed to the emergence of numerous phenotypic traits that distinguish modern humans from their closest extinct relatives.

### Human-specific candidate regulatory variants are enriched in large deserts of archaic introgression

If some of the candidate regulatory variants identified here contributed to phenotypic divergence between modern humans and archaic hominins, they may also have reduced the fitness of hybrid individuals following admixture. Under this scenario, genomic regions carrying such variants would be expected to show reduced levels of archaic ancestry due to purifying selection acting against introgressed haplotypes. The enrichment for genes with autosomal dominant phenotypes (Table 1) linked to the human-specific candidate regulatory variants supports this hypothesis.

Previous studies have shown that the removal of archaic DNA from the modern human genome was particularly pronounced in functionally important regions and that much of this purging occurred rapidly during the generations immediately following admixture (29-37). Under such conditions, selection acts on entire introgressed haplotypes before recombination has sufficient time to break them down into smaller fragments. As a consequence, strongly deleterious archaic alleles can generate extended regions depleted of archaic ancestry surrounding the selected locus.

To investigate if our human-specific variants show evidence for this possibility, we intersected them with regions of the human genome depleted of Neanderthal and Denisovan ancestry (38, 39). We found that 119 candidate variants out of the 404 candidate regulatory variants occur within 86 regions lacking archaic introgression (Supplementary Table 13).

These regions varied substantially in size, ranging from 6,138 bp to more than 8,909,000 bb, with a mean length of 591,320 bp (Figure 4).

**Figure 4.**
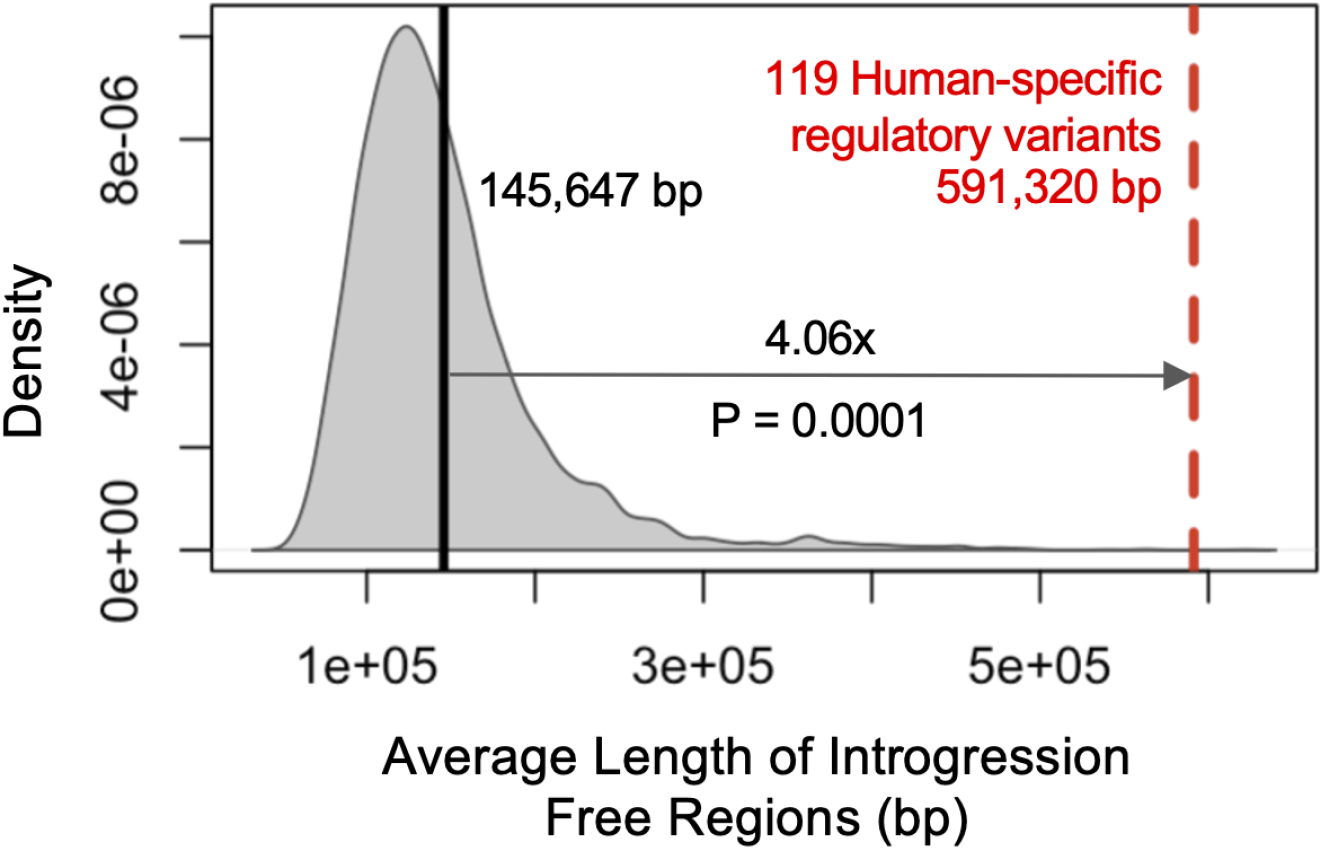
Human-specific regulatory variants are enriched in long regions depleted of archaic introgression. Human-specific regulatory variants are enriched in introgression-depleted regions that are significantly longer than expected by chance (4.06x, P = 0.0001; Supplementary Table 13).

To determine whether our variants fall in larger depleted regions than expected, which would be consistent with selection against introgressed archaic alleles, we compared the observed length distribution with null distributions generated from randomly sampled sets of regions lacking introgression. The average size of the regions free of archaic introgression containing candidate human-specific regulatory variants was more than four times greater than expected (P = 0.0001), indicating that these variants preferentially occur within larger regions lacking archaic ancestry (Figure 4B).

This is consistent with the hypothesis that some of these loci were targets of purifying selection following admixture between modern humans and archaic hominins. This is particularly intriguing given that the variants were identified based on their potential effects on traits that distinguish modern humans from Neanderthals.

## Discussion

A central challenge in human evolutionary genomics is moving from catalogs of genetic variants with evidence of selection to an understanding of the phenotypes they influence. Ancient DNA has revealed thousands of genomic changes distinguishing modern humans from archaic hominins, but inferring organism-level effects of these variants has proven difficult. Here we contribute to this effort with a framework for identifying human-specific variants with strong regulatory potential and connecting them to candidate phenotypes by requiring convergent evidence across independent functional layers: disruption of a TFBS, location within an experimentally annotated cis-regulatory element, and a phenotype association shared between the target gene and the TF gene. The presence or absence of a predicted motif alone does not provide enough evidence for biological relevance. This multi-layered approach allows us to move from a long list of candidate variants to a prioritized set of testable genotype-phenotype hypotheses with direct relevance to human-specific traits.

We identified 404 human-specific candidate regulatory variants linked to 387 genes. Several independent lines of evidence validate the biological function and evolutionary relevance of this candidate set. Human-specific variants predicted to disrupt or create a TFBS in known cCREs were significantly enriched among regions differentially methylated between modern humans and archaics. Human-specific variants validated to alter gene regulation in an MPRA assay were also enriched among our candidate set. Finally, the genes linked to the human-specific variants are also significantly enriched for traits known to be different among humans and archaics. Specifically, we identified 128 variant and 155 candidate genes functionally involved in at least 58 traits that are known to differentiate humans and archaics. The candidate variants are also enriched in long regions depleted of archaic introgression, consistent with the idea that they contributed to phenotypic changes and that the corresponding archaic alleles were rapidly purged by purifying selection after admixture. The convergence of evolutionary, methylation, expression, and phenotype-association-based evidence around these variants strengthens the case that they produce functional regulatory divergence.

The phenotypes associated with these variants confirm previously proposed differences known from the fossil record. However, we are also able to extend beyond preserved traits and propose candidate phenotypic differences that cannot be directly observed in skeletal remains. For example, we find enrichment for differences in brain and behavioral traits such as autism and the hypothalamus, reproductive traits like cryptorchidism, facial traits that do not fossilize like the lips and for speech and language abilities. Several of these are consistent with earlier suggestions of soft tissue divergence between modern humans and archaics. Our results provide independent regulatory support for these earlier hypotheses, and they highlight specific variants with mechanistic functional hypotheses about their effects on gene regulation. They additionally point to previously unreported candidate traits, such as those related to reproductive anatomy, that merit further investigation.

The observation that candidate human-specific regulatory variants are preferentially located within long genomic regions depleted of archaic introgression raises the possibility that at least some of these regulatory changes contributed not only to phenotypic divergence, but also led to reduced fitness in human-archaic hybrids carrying the ancestral allele due to introgression. This is consistent with a role for regulatory incompatibilities and the partial reproductive isolation between these lineages. If true, this would place at least part of the genetic basis of hybrid incompatibility in the same class of regulatory changes responsible for morphological divergence, linking the evolution of phenotype divergence directly to the process of lineage separation.

Beyond the specific candidates highlighted here, our results provide a resource to guide future functional work. Researchers can use the prioritized variant-gene-phenotype combinations to select candidates most likely to have measurable regulatory consequences on phenotypes of interest, focusing experimental effort where converging computational and functional evidence already exists. Moreover, the approach itself is not specific to the comparison between modern humans and archaic hominins: the same pipeline could be extended to other members of the genus *Homo* as additional archaic genomes become available, or applied more broadly to any pair of species with characterized TFBS annotations and phenotype ontologies, such as mouse or chicken, to study the regulatory basis of trait divergence across the tree of life.

## Conclusion

By integrating human-specific genetic variants, transcription factor binding site motifs, and phenotype ontologies, we identified 404 candidate regulatory variants implicating 387 genes and 586 traits that may distinguish modern humans from Neanderthals and Denisovans. These candidates are independently supported by patterns of differential DNA methylation, by experimentally measured changes in gene regulatory activity, and alignment with known traits differentiating humans and archaics. The discovered gene-to-phenotype annotations extend beyond traits preserved in the fossil record to point to previously suggested soft-tissue phenotypes such as lips, brain, testes and language differences. Their enrichment within unusually large regions depleted of archaic introgression further suggests that some of these regulatory changes may have contributed to reduced fitness in human-archaic hybrids. Together, these results provide a prioritized set of variants, mechanistic regulatory hypotheses, and a generalizable framework for connecting human-specific regulatory variation to the phenotypes that define our species, with potential application to other *Homo* lineages and other species with well characterized regulatory annotations.

## Supporting information

https://docs.google.com/spreadsheets/d/1xMaJcX0Xq92T89GGD4RwN_LOdrrlfs9as4-qxeNB-Lw/edit?gid=314984489#gid=314984489

## Methods

### Human-specific variants

Human-specific variants were obtained from two studies that compared thousands of individuals from the 1000 Genomes Project with three or four high-coverage Neanderthal genomes, one Denisovan genome, and several individuals from four great ape species (1, 3). Both studies differ in their criterion to determine if a variant is nearly fixed. From the first study, we retained only those variants that were fixed for one allele in modern humans and fixed for the alternative allele in archaic hominins. From the second study, we retained all variants because they have already undergone functional characterization, making them valuable candidates for downstream analyses. We lifted these variants over from human reference genome hg19 to the human reference genome hg38 with a liftover tool from UCSC (40). After merging both datasets, removing redundant entries and variants with conflicting human-archaic allele calls, we obtained a set of 16,883 variants.

### GPN-MSA DNA language model

Scores for all variants were taken from pre-computed GPN-MSA scores (16). GPN-MSA (genomic pretrained network with multiple-sequence alignment) is a DNA language model that leverages whole-genome multiple sequence alignments from 100 vertebrate species to score the tolerance for each possible allele at single-nucleotide resolution for the human genome. Of our human-specific variant set, 16,639 (98.6%) could be scored by GPN-MSA, allowing a direct comparison between the human-specific and archaic alleles.

### Identification of cCREs

Candidate cCREs for hg38 were downloaded from the ENCODE Registry of cis-Regulatory Elements (cCREs) via SCREEN (18). The set is meant to be a genome-wide cell-type agnostic catalog of candidate regulatory elements defined from chromatin accessibility, histone modifications, and transcription factor binding data.

### Identification of changes in TFBS motifs

Changes in TFBSs affinity for the human-specific variants were predicted with FABIAN-variant 2026, a tool that evaluates the effect of a DNA variant on TF binding by combining two types of motif model: position weight matrices (PWMs) and transcription factor flexible models (TFFMs, an extension of hidden Markov models that also capture dependencies between adjacent positions) (17). For each variant, FABIAN-variant compares predicted binding affinity for the reference and alternative alleles across all available models for a given transcription factor and returns a combined score reflecting predicted loss or gain of binding. We retained only variant–TF predictions with a combined score ≥0.9 or ≤−0.9 in at least one of the models tested. Each model assigns values from -1 (strong evidence of disrupting a specific TFBS motif) to 1 (strong evidence of creating a motif). This retains <0.05% of the tested models.

### Target gene annotations

For gene annotation, we assigned each human-specific variant to its closest gene when the same target gene was identified by both GREAT (41) and FUMA (42). GREAT mapping is based on proximity, and FUMA integrates positional mapping, eQTL mapping, and chromatin interaction mapping to identify target genes.

### Known human-specific skeletal traits

Traits known to differ between humans and archaics based on the fossil record were taken from Gokhman et al. (2019) (19), who classified 361 traits based on HPO annotations. Using the current version of the HPO gene-to-phenotype annotation file (July 2026) (20), only 341 of these traits still had associated gene annotations. We restricted our analysis to these 341 traits. Organ-level enrichment was assessed using Gene ORGANizer (22) since it has been previously used to predict differences in soft tissues among humans, Neanderthals, Denisovans, and chimpanzees (3, 23). For the organ enrichment among the target genes we used the background of all genes with HPO annotations in geneORGANizer. For the enrichment across TFs, we used all known TFs in the Fabian-Variant database to correct for a possible bias given the importance of TFs in gene-to-phenotype associations. Candidate phenotypes were also identified through enrichment analysis of HPO terms using Enrichr (21).

### Differences in methylation and regulatory activity between humans and archaics

Differences between hominin groups have previously been reported in methylation maps (DMRs) and in ability to drive transcription in MPRAs in neural progenitor cells, stem cells, and osteoblasts. Methylation data were taken from Gokhman et al. (2020) (23), and MPRA activity data from Weiss et al. (2021) (3) . For permutation analyses involving DMRs, variants on the X chromosome were excluded, as no DMR maps are available for the sex chromosomes.

### Regions lacking archaic introgression

Regions depleted of Neanderthal introgression were obtained from Chen et al. (2026) (38) and Liang et al. (2025) (39). Regions depleted of Denisovan introgression were taken from Vernot et al. (2016) (36). Human-specific variants outside any of the detected introgressed segments were classified as variants in regions free of introgression. Depending on the method, each variant could be assigned to several segments varying in length. For each variant we kept the shortest segment. 119 of the 404 candidate variants are in 86 regions lacking introgression (74 in the autosomes and 12 in the X chromosome). To test if the candidate variants are in larger regions free of archaic ancestry than expected, we conducted 10000 permutations. Given that the longest deserts of archaic introgression are in the X chromosome (31-38), we maintained the observed pattern (74 regions in the autosomes and 12 in the chromosome X) in each permutation.

### Enrichment for known divergent skeletal traits

Across all variants for which the target gene and transcription factor (TF) were associated with the same HPO trait, we identified 261 target genes and 149 TFs, forming multiple target–TF combinations, with some genes and TFs occurring only once and others occurring multiple times. These combinations are collectively associated with 58 skeletal traits known to differ between modern humans and archaic hominins. To assess whether this overlap was greater than expected by chance, we performed 10,000 permutations by randomly sampling 261 protein-coding genes from all protein-coding genes with HPO annotations and 148 TFs from all TFs with HPO annotations. For each permutation, genes and TFs were sampled with the same multiplicity structure observed in the empirical data, such that genes and TFs appearing once or multiple times in the observed sets were represented with the corresponding frequencies in the permuted sets. The two resulting lists were independently shuffled and paired to generate randomized target–TF combinations. For each permutation, we counted the number of skeletal traits known to differ between modern humans and archaic hominins that are associated with at least one of these randomized combinations. The observed data showed a significantly higher number of skeletal traits than the values obtained across the 10,000 permutations (P < 0.0002), indicating that the observed target–TF combinations are enriched for regulatory relationships associated with skeletal traits that differ between modern humans and archaic hominins.

